# Interferon-γ and IL-27 positively regulate type 1 regulatory T-cell development during adaptive tolerance

**DOI:** 10.1101/2024.06.21.598825

**Authors:** David A.J. Lecky, Lozan Sheriff, Sophie T. Rouvray, Lorna S. George, Rebecca A. Drummond, David C. Wraith, David Bending

## Abstract

Strong T-cell receptor (TCR) and IL-27 signalling influence type-1 regulatory (Tr1) T-cell development but whether other signals determine their differentiation is unclear. Utilising Tg4 TCR transgenic mice we established a model for rapid Tr1 cell induction. A single high dose of [4Y]-MBP peptide drove the differentiation of *Il10*^+^ T-cells with *bona fide* Tr1 cell protein and mRNA signatures. Kinetic transcriptional analysis revealed that the Tr1 cell module was transient and preceded by a burst of *Ifng* transcription in CD4+ T-cells. Neutralisation of IFNγ reduced Tr1 cell frequency and strong TCR signalling markers, which was correlated with reduced macrophage activation. Antibody depletion experiments inferred that T-cells – but not NK cells – provided the relevant source of IFNγ. Additionally, we show that blocking IL-27 in combination with IFNγ neutralisation additively reduced Tr1 cell frequency *in vivo*. These findings reveal that during strong tolerogenic TCR signalling IFN-γ has a non-redundant regulatory role in augmenting the differentiation of Tr1 cells *in vivo*.

## Introduction

Interleukin(IL)-10 is an immunoregulatory cytokine produced by multiple immune cell types but has a key function in the regulation of adaptive immunity ^1^. IL-10 can inhibit antigen presentation, cell proliferation and pro-inflammatory cytokines ^2,3^, including directly countering interferon(IFN)γ through inhibition of IFNγ-induced genes, including Major Histocompatibility Complex (MHC) Class II ^4,5^. MHC II presents peptide fragments to T-cell Receptor (TCR) complexes on CD4^+^ T-cells, to promote T-cell activation. IL-10 inhibits antigen specific proliferative T-cell responses, by reducing the antigen presentation capacity of Dendritic Cells (DCs), monocytes, macrophages, and other Antigen Presenting Cells (APCs) through MHCII downregulation ^6^, limiting co-stimulation ^7^ and production of inflammatory cytokines like IL-12 ^8^. Reduction in constitutive and IFNγ-induced MHCII expression in monocytes leads to inhibition of antigen-specific T-cell responses, partially contributing to tolerance of presented antigens, including tumour-derived antigens.

Amongst CD4^+^ T-cells, major producers of IL-10 are FoxP3^+^ Treg cells and Type 1 regulatory T (Tr1) cells ^9^. In Tr1 cells, IL-10 is co-expressed with Lymphocyte-activation gene 3 (LAG3) and can be induced by the cytokine IL-27 ^10^, which can in turn activate the transcription factor c-Maf to promote Tr1 cell differentiation ^11^. The Tr1 cell associated markers T-cell immunoreceptor with immunoglobulin and ITIM domain (TIGIT) and LAG3 bind CD155 and MHCII respectively, interfering with optimal TCR transduction and immunological synapse formation ^12,13^. Egr-2 ^14^ in combination with Blimp-1 ^15^ is required to mediate IL-10 production in IL-27 activated murine CD4^+^ T-cells.

IL-10 expression has also been linked to the relative strength of TCR signalling ^16^, and repeated dosing of Tg4 TCR transgenic mice with a modified self-peptide leads to the progressive emergence of FoxP3^−^ Tr1-like IL-10^+^ T-cells ^17^. We recently developed an accelerated adaptive tolerance model ^18^ that induces expression of *Il10*-enhanced Green Fluorescent Protein (eGFP) within 24 hr of primary *in vivo* immunisation with a high dose of self-antigen ^19^. *Il10*-eGFP^+^ T-cells arose from T-cells experiencing the highest levels of TCR signal strength *in vivo* and expressed higher levels of LAG3 and TIGIT ^19^. Despite uniform activation of T-cells in the accelerated adaptive tolerance model, as evidenced by activation of NFAT-driven *Nr4a3* transcription, only 15-25% of activated T-cells go on to transcribe *Il10*. This finding suggests that high antigen concentrations alone are not sufficient for promoting *Il10* transcription and that other factors and cellular interactions within the local milieu play important roles. Here, we established a model for the rapid induction of Tr1 cells *in vivo*. Our system revealed a non-redundant regulatory role for IFNγ in combination with IL-27 to promote Tr1 cell differentiation under conditions of strong tolerogenic TCR signalling.

## Results

### Kinetics of *de novo Il10* transcription in response to strong tolerogenic TCR signalling

We previously reported that strong TCR signalling drives *Il10* transcription ^19^. To further investigate the development of *Il10*^+^ T-cell populations and their relationship to Tr1 cells in this model we performed tolerogenic immunisation of Tg4 *Nr4a3*-Tocky *Il10*-eGFP transgenic mice with a single dose of [4Y] Myelin Basic Protein (MBP) peptide. Firstly, we established baseline expression of downstream TCR signalling (*Nr4a3*-Timer, hereafter referred to as “Timer”) and *Il10*-eGFP reporters under a low, medium, and high (0.8, 8 and 80 μg) dose of [4Y]-MBP peptide at 24 hr. As *Nr4a3*-Timer expression is downstream of TCR engagement and NFAT signalling ^20^, we use it as a proxy marker for TCR specific activation. Timer^+^ population positively correlated with [4Y]-MBP peptide dose (**Figure 1A**) and established that *Il10*-eGFP was almost exclusively found within the Timer^+^ fraction (i.e., recently TCR signalled, **Figure 1B**). In the Timer^+^ population, LAG3 (**Figure 1C**) and TIGIT (**Figure 1D**) positively correlated with peptide dose only within the Timer^+^ fraction, highlighting the importance of TCR signalling strength for induction of IL-10, LAG3 and TIGIT in this model. Furthermore, we also observed dose-dependent increases in the levels of LAG3 (**Figure 1E**) and TIGIT (**Figure 1F**) within *Il10*-eGFP^+^ T-cells.

**Figure 1:**
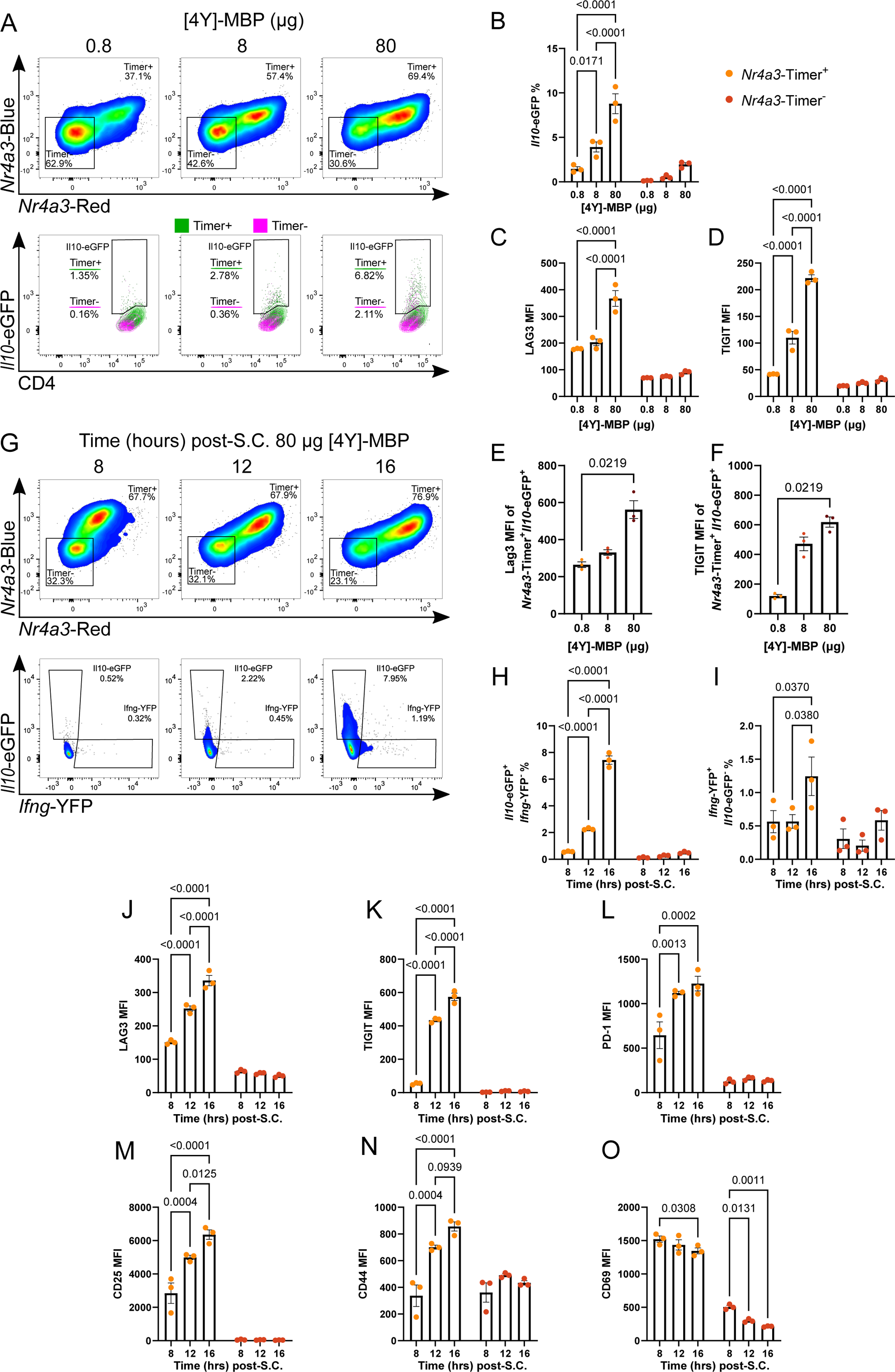
Kinetics of *de novo Il10* transcription in response to strong tolerogenic TCR signalling. Tg4 *Nr4a3*-Tocky *Il10*-eGFP mice were immunised with 0.8, 8, or 80 μg [4Y]-MBP in 200 μL PBS s.c. and spleens were harvested at 24 hr. Representative flow plots **(A)** showing CD4^+^ TCRvβ8.1/8.2^+^ according to *Nr4a3*-Timer expression above and the *Il10*-eGFP^+^ of *Nr4a3*-Timer^+^ below. Summary of *Il10*-eGFP frequency **(B)**, LAG3 MFI **(C)**, and TIGIT MFI **(D)** in *Nr4a3*-Timer^±^. Summary of LAG3 **(E)** and TIGIT MFI **(F)** in *Il10*-eGFP^+^ *Nr4a3*-Timer^+^. Tg4 *Nr4a3*-Tocky *Il10*-eGFP *Ifng*-YFP mice were administered 80 μg [4Y]-MBP in 200 μL PBS s.c. and spleens were harvested at 8, 12 or 16 hr. Representative flow plots **(G)** showing CD4^+^ TCRvβ8.1/8.2^+^ *Nr4a3*-Timer expression above and below *Il10*-eGFP against *Ifng*-YFP of *Nr4a3*-Timer^+^ below. Summary of *Il10*-eGFP^+^ *Ifng*-YFP^−^ **(H)** and *Ifng*-YFP^+^ *Il10*-eGFP^−^ **(I)** frequency, LAG3 **(J)**, TIGIT **(K)**, PD-1 **(L)**, CD25 **(M)**, CD44 **(N)** and CD69 **(O)** MFI in *Nr4a3*-Timer^±^. **(B-F, H-O)** bars represent median with interquartile range or mean+/−SEM. Statistical analysis by two-way ANOVA with Sidak’s multiple comparisons test (**B-D**, **H-O**) or Kruskal Wallis test with Dunn’s multiple comparisons. N = 3 per treatment or time.

IL-10 expression has been previously observed in the Tg4 tolerance model to arise from IFNγ producing Th1 cells that have undergone repeat stimulation ^8^. To assess the relationship between the development of *Il10*-eGFP and IFNγ production, we performed kinetic analysis at 8, 12 and 16 hr following high dose of [4Y]-MBP (80 μg) stimulation of Tg4 *Nr4a3*-Tocky *Il10*-eGFP *Ifng*-YFP transgenic mice. Timer expression shifts from mostly “blue^+^” to mostly “blue^+^ red^+^” over the time course (**Figure 1G**, top panel), representing a shift from recently activated (< 4 hr) to persistent activation (> 4 hr, < 24 hr). *Il10*-eGFP and *Ifng*-YFP are also increasingly expressed within this Timer^+^ population and predominantly arise independently of each other (**Figure 1G**, bottom panel). *Il10*-eGFP significantly and steadily increases between 8 and 16 hr in the Timer^+^ population (**Figure 1H**), with no significant increases in *Il10*-eGFP for the Timer^−^ fraction. Within CD4^+^ T-cells, *Ifng*-YFP is expressed in both Timer^+^ and Timer^−^, with peak detection at 16 hr (**Figure 1I**). In the Timer^+^ cells, LAG3, TIGIT, PD-1, CD25, and CD44 (**Figures 1J-N**, respectively) show increasing expression from 8 hr to 16 hr, only within the Timer^+^ fractions. Notably, CD69 does not follow this pattern, showing decreasing expression in both Timer positive and negative (**Figure 1O**) implying this marker becomes decoupled from TCR signalling in this model. These findings demonstrate that CD4^+^ T-cells can rapidly transcribe *Il10* and increase expression of Tr1 markers in a TCR signal strength and time dependent fashion during the first 24 hours of immunisation.

### Rapidly induced *Il10* expressing T-cells are Tr1-like cells and have a transcriptionally delayed programme

To confirm that the *Il10*-eGFP population arose from a non FoxP3^+^ Treg precursor, we sorted CD4^+^ Timer^+^ T-cells by *Il10*-eGFP positivity or negativity 24 hrs after immunisation (**Figure 2A** and **2B**). Intracellular FoxP3 staining of sorted CD4^+^ *Il10*-eGFP^+/−^ (**Figure 2C**) shows that *Il10*-eGFP^+^ CD4 T-cells are largely FoxP3^−^ (∼98 %), whereas it is the larger proportion of the *Il10*-eGFP^−^ that are enriched for FoxP3 (∼8 %, **Figure 2D**). FoxP3 expression, therefore, is not associated with CD4^+^ T-cell *Il10*-eGFP expression in our model. We similarly sorted CD4^+^ *Il10*-eGFP^+^ and CD4^+^ *Il10*-eGFP^−^ T-cells to undergo mRNA-seq analysis at 24 hrs (**Figure 2E**). Heatmap analysis revealed that the *Il10-*eGFP^+^ subset was enriched with transcripts for hallmark signatures of Tr1 cells, including *Maf*, *Prdm1* (encoding Blimp1)*, Il10* and *Ctla4*. In addition, they also showed enhanced expression of *Il12rb2* and *Ifngr1* both of which encode receptors important in the generation of Th1 responses.

**Figure 2:**
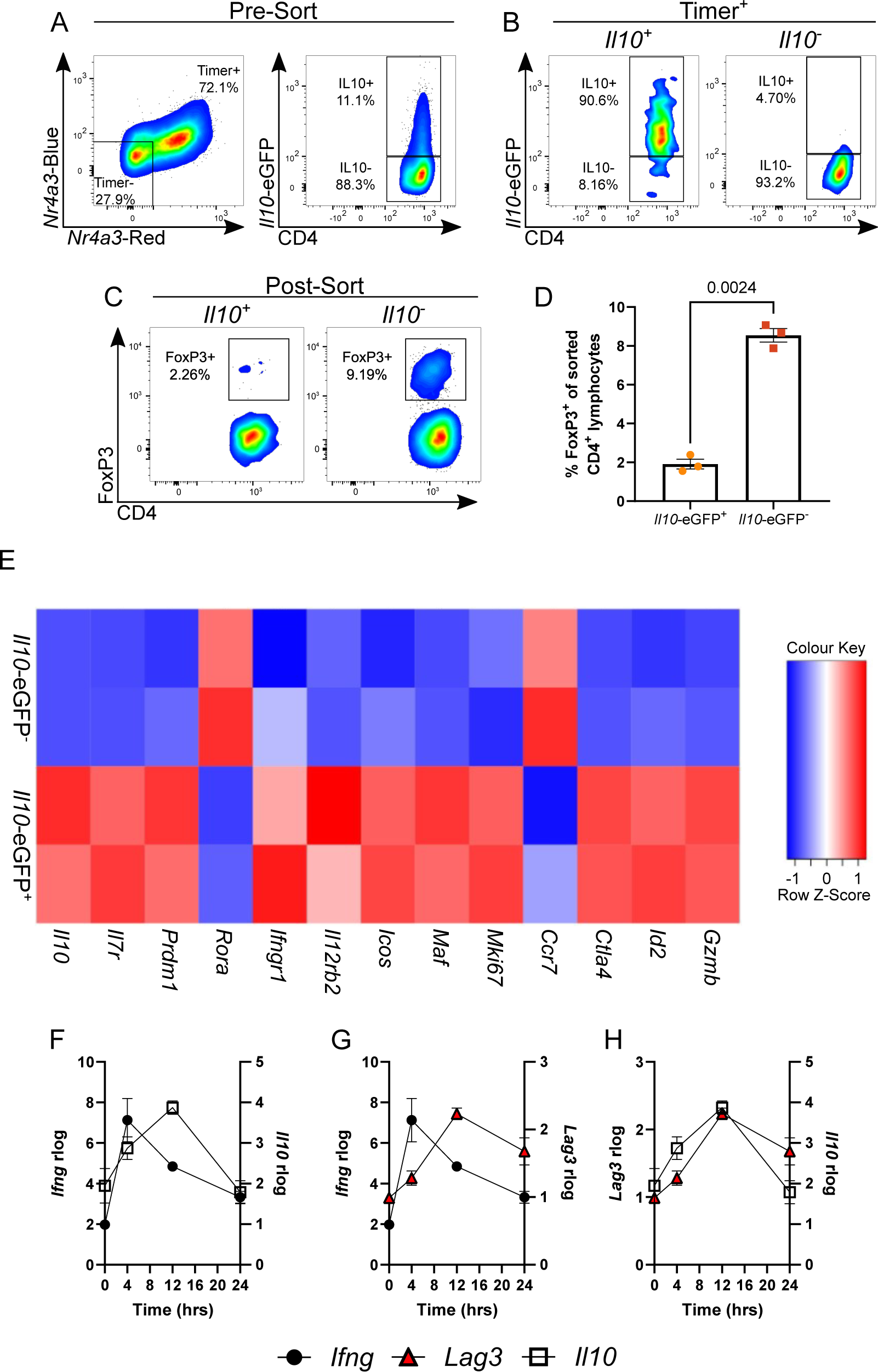
Rapidly induced *Il10* expressing T-cells are Tr1-like cells and have a transcriptionally delayed programme. Tg4 *Nr4a3*-Tocky *Il10*-eGFP mice were administered 4 mg/kg [4Y]-MBP in 200 μL PBS s.c. and spleens were harvested at 24 hr. Representative flow plots of CD4^+^ TCRvβ8.1/8.2^+^ pre-sort showing *Nr4a3*-Timer and *Il10*-eGFP expression **(A)**, post-sort *Il10*-eGFP in *Nr4a3*-Timer^+^ **(B)**, and post-sort FoxP3 expression in CD4^+^ TCRvβ8.1/8.2^+^ *Nr4a3*-Timer^+^ *Il10*-eGFP^±^ **(C)**. Summary of FoxP3 frequency by *Il10*-eGFP^±^ **(D)**. Transcriptome analysis of CD4^+^ *Nr4a3*-Timer^+^ *Il10*-eGFP^±^ as a heatmap of curated Tr1 DEGs between conditions at 24 hr **(E)**. Compared temporal expression rlog transcripts from GEO: GSE165817 (Elliot et al. ^19^) for *Ifng* against *Il10* **(F)**, *Ifng* against *Lag3* **(G)**, and *Lag3* against *Il10* **(H)** from Tg4 *Nr4a3*-Timer *Il10*-eGFP mice administered with 80 μg [4Y]-MBP in 200 μL PBS s.c. and mRNA transcripts depicted at 4, 12 and 24 hr. Bars represent mean+/− SEM, statistical analysis by paired t test. N = 3 per group.

To understand the kinetics of development of the Tr1 transcriptional programme, we re-analysed a past RNA-seq time course experiment performed on CD4^+^ T-cells that had received 0 μg (control), 0.8 μg or 80 μg [4Y]-MBP for up to 24 hrs *in vivo* ^19^. Here we focused on markers associated with TCR signalling, Th1 cells and Tr1 phenotype (**Supplementary Figure 1**). Our analysis showed that strong tolerogenic stimulation led to the rapid activation of the early activation genes *Nr4a3* and *Tnf* which was also linked to a rapid transcriptional burst of *Tbx21*, *Ifng* and *Il2* (Th1-type signature) at 4 hours. The Tr1 module (comprising *Il10, Tigit, Lag3, Maf3, Nfil3, Prdm1*) *appeared* delayed and arising more uniformly at 12 hours and was only visible in the condition of strong TCR signalling. Given the potential relevance of the early type-1 immune signature, we compared the transcriptional changes at 4, 12 and 24 hr timepoints post-peptide administration for *Ifng, Il10* and *Lag3* (**Figure 2F-H**). Analysis confirmed that the *Ifng* transcript peak preceded the *Il10* transcriptional peak by around 8 hrs, with *Ifng* peaking at 4 hrs, and *Il10* at 12 hr (**Figure 2F**). *Lag3* transcriptional peak also showed a similar relationship to *Ifng* as *Il10* (**Figure 2G**), which was clearly observed when *Il10* and *Lag3* transcriptional kinetics were compared head to head (**Figure 2H**). Finding that *Ifng* transcriptionally preceded the emergence of Tr1 cells our accelerated adaptive tolerance model, we hypothesised that it may play a role in regulating Tr1 cell development.

### Interferon-gamma positively regulates Tr1 cell differentiation

Having established that *Il10*-eGFP is expressed by rapidly induced Tr1 cells, and the *Il10* transcript is preceded by *Ifng* transcript, we administered αIFNγ antibody before weight-normalised immunisation of Tg4 *Nr4a3*-Tocky *Il10*-eGFP *Ifng-*YFP *mice* with 4 mg/kg of [4Y]-MBP peptide. At 24 hr, splenic CD4^+^ T-cell responses were analysed by flow cytometry (**Figure 3A**). Analysis of Tg4 *Nr4a3*-Timer^+^ T-cells revealed that the frequency of *Il10*-eGFP (**Figure 3B**) but not *Ifng*-YFP (**Figure 3C**) expressers was significantly reduced by αIFNγ treatment. αIFNγ treatment did not alter CD69, TIGIT or LAG3 expression (**Figure 3D-F**) but did significantly reduce ICOS and GITR (**Figure 3G&H**), which we have previously shown to be markers of TCR signal strength in this model ^19^.

**Figure 3:**
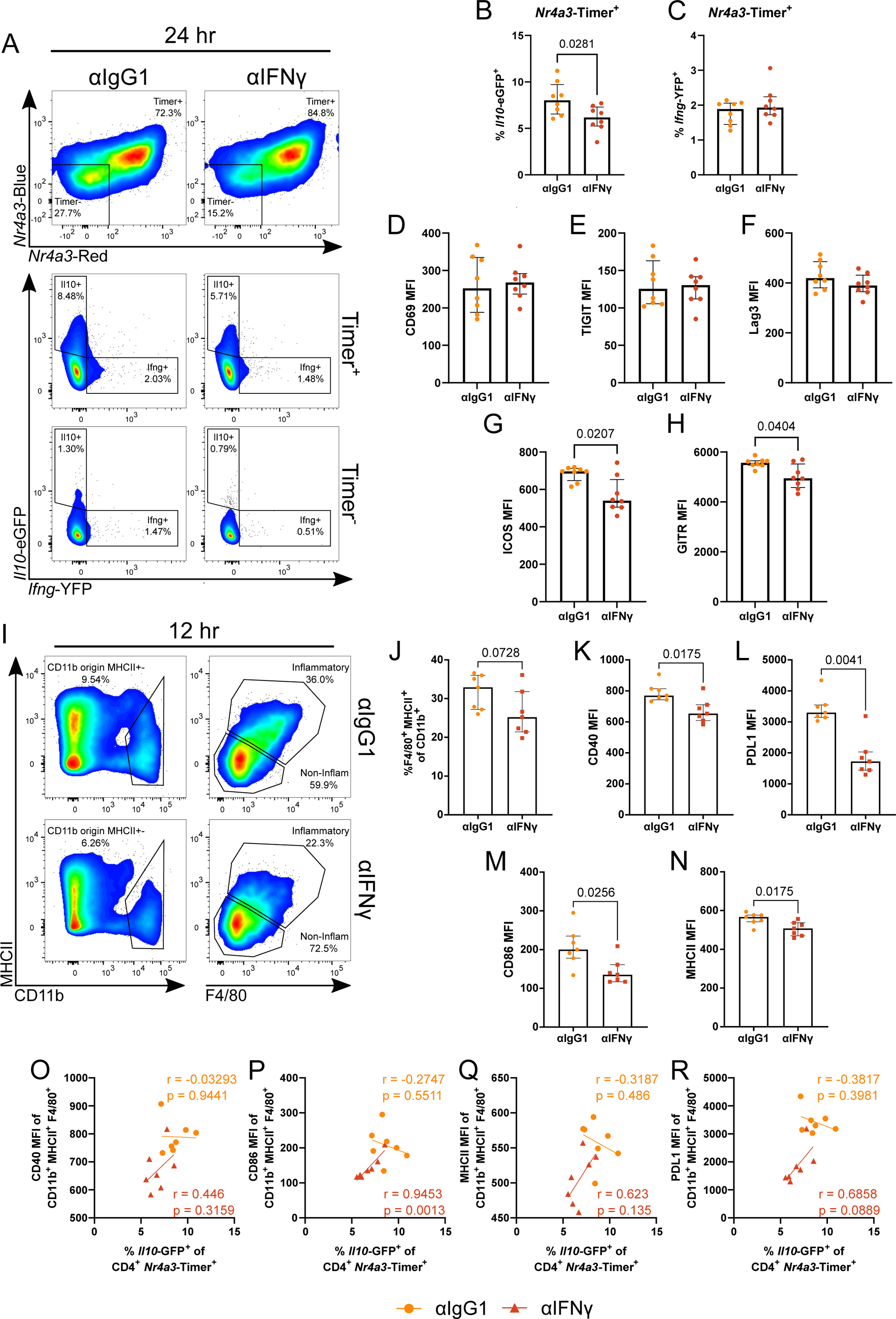
Interferon-gamma positively regulates Tr1 cell differentiation. Tg4 *Nr4a3*-Tocky *Il10*-eGFP *Ifng*-YFP mice were immunised with 4 mg/kg [4Y]-MBP in PBS s.c. and 1 mg αIFNγ or αIgG1 isotype in 200 μL PBS i.p. and spleens were harvested 24 hours later. Representative flow plots at 24 hr **(A)** showing CD4^+^ TCRvβ8.1/8.2^+^ according to *Nr4a3*-Timer expression above and the *Il10*-eGFP against *Ifng*-YFP of *Nr4a3*-Timer^+^ below. Summary of *Il10*-eGFP **(B)** and *Ifng*-YFP **(C)** frequency in *Nr4a3*-Timer^+^, and CD69 **(D)**, TIGIT **(E)**, LAG3 **(F)**, ICOS **(G)**, and GITR **(H)** MFI. Tg4 *Nr4a3*-Tocky *Il10*-eGFP *Ifng*-YFP mice were immunised with 4 mg/kg [4Y]-MBP in PBS s.c. and 1 mg αIFNγ or αIgG1 isotype in 200 μL PBS i.p. and spleens were harvested at 12 hours **(I)** showing CD11b against MHCII to the left, and from CD11b^+^, F4/80 against MHCII to the right. Summary of F4/80^+^ MHCII^+^ (“Inflammatory” myeloid) frequency **(J)** CD40 **(K)**, PD-L1 **(L)**, CD86 **(M)**, and MHCII **(N)** MFI. Linear regression (Pearson’s correlation) of CD4^+^ *Nr4a3*-Timer^+^ *Il10*-eGFP^+^ frequency against CD40 **(O)**, CD86 **(P)**, MHCII **(Q)** and PD-L1 **(R)** MFI in the inflammatory myeloid compartment. **(B-H, I-N)** bars represent median with interquartile range. Statistical analysis by Mann-Whitney U test. N = 8 **(A-I)**, N = 7 **(J-S)** per treatment.

Given the changes in *Il10* and markers of TCR signal strength, we hypothesised that IFNγ may have a role in augmenting TCR signal strength and hence the induction of Tr1 cells in our model. We performed analysis of splenic DC, B cell and macrophage subsets. Analysis revealed a general reduction in CD19^−^ MHC Class II^+^ cells within the splenic environment but had no impact on DC frequency but did reduce DC levels of PD-L1 (**Supplementary Figure 2**). Analysis of splenic macrophages, identified as MHC Class II^+^ F4/80^+^ cells from a CD11b^+^ origin^21^ (**Figure 3I**) revealed a strong trend for a reduction in this population (**Figure 3J**) and that these splenic macrophages all had significantly reduced levels of CD40 (**Figure 3K**), PD-L1 (**Figure 3L**), CD86 (**Figure 3M**) and MHC Class II (**Figure 3N**). Correlation analysis of the frequency of *Il10-*eGFP^+^ T-cells in comparison to macrophage activation status highlighted that within αIFNγ treated groups, all showed positive correlations (**Figure 3O-R**) with CD86 significantly and strongly correlating with Tr1 cell frequency. These data demonstrate that IFNγ plays a positive regulatory role in enhancing Tr1 cell development, which is correlated with reduced levels of TCR signal strength and myeloid cell activation.

### NK and T-cells are major sources of IFNγ but NKs are redundant for Tr1 cell induction

Next, we wanted to determine the major splenic sources of IFNγ *in vivo*. Following immunisation of Tg4 *Nr4a3*-Tocky *Il10*-eGFP *Ifng-*YFP mice and treatment with isotype or αIFNγ, we further analysed the *Ifng*-YFP^+^ compartments (**Figure 4A**). NK1.1^+^ TCR^−^ cells (**Figure 4A**) comprised the majority of the *Ifng-*YFP producers, with the remainder being largely T-cell derived. Interestingly, total splenic frequency of *Ifng*-YFP^+^ cells was unaffected by αIFNγ treatment but did drive a significant increase in *Ifng*-YFP levels in NK1.1^+^ cells (**Figure 4B-D**).

**Figure 4:**
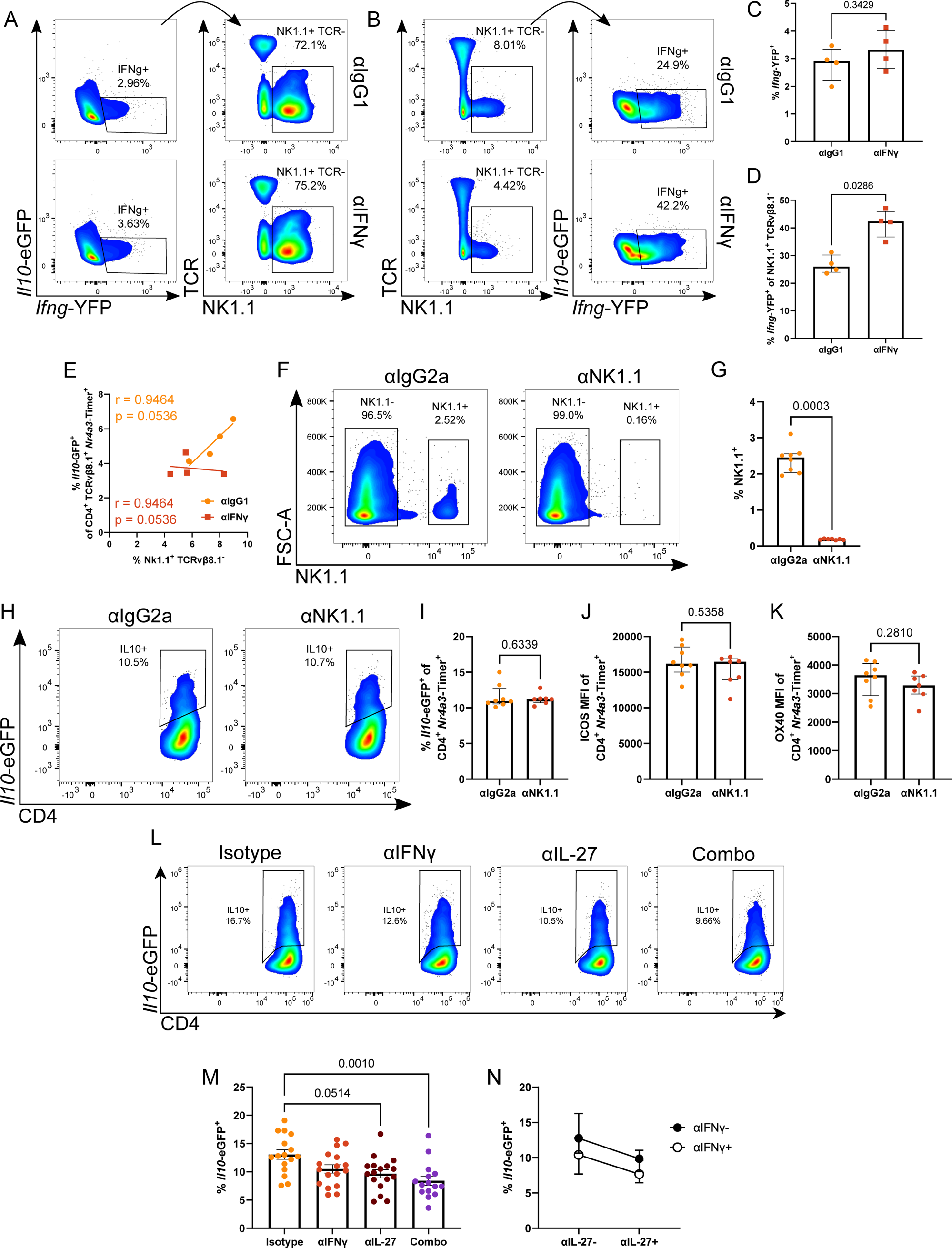
IFNγ and IL-27 additively regulate Tr1 cell development. Tg4 *Nr4a3*-Tocky *Il10*-eGFP *Ifng*-YFP mice were administered 4 mg/kg [4Y]-MBP in PBS s.c. and 1 mg αIFNγ or αIgG1 isotype in 200 μL PBS i.p. and spleens were harvested at 24 hr. Representative flow plots showing *Il10*-eGFP against *Ifng*-YFP above and *Ifng*-YFP^+^-derived TCRvβ8.1/8.2 against NK1.1 below **(A)**, and TCRvβ8.1/8.2 against NK1.1 above and NK1.1^+^-derived *Il10*-eGFP against *Ifng*-YFP below **(B)**. Summary of *Ifng*-YFP^+^ frequency **(C)** from **(A)** and *Ifng*-YFP^+^ frequency from NK1.1^+^ **(D)** from **(B)**. Linear regression (pearson’s correlation) of CD4^+^ *Nr4a3*-Timer^+^ *Il10*-eGFP^+^ against NK1.1^+^ TCRvβ8.1/8.2^−^ **(E)**. Tg4 *Nr4a3*-Tocky *Il10*-eGFP mice were given 200 μg αNK1.1 or αIgG2a isotype in 200 μL PBS i.p and 48 hrs later were immunised with 4 mg/kg [4Y]-MBP in PBS s.c. and. and spleens were harvested 24 hr later. Representative spectral cytometry plots showing NK1.1 against FSC-A **(F)**. Summary of NK1.1^+^ **(G)** from **(F)**. Representative spectral cytometry plots at 24 hr showing *Il10*-eGFP against CD4 **(H)**. Summary of *Il10*-eGFP^+^ frequency **(I)** and ICOS **(J)** and OX40 **(K)** MFI of CD4^+^ *Nr4a3*-Timer^+^ T-cells. Tg4 *Nr4a3*-Tocky *Il10*-eGFP mice were administered 4 mg/kg [4Y]-MBP in PBS s.c. and 0.5 mg αIFNγ, αIL-27, a combination of both or a mixed IgG1 and αIgG2a isotype in 200 μL PBS i.p. and spleens were harvested 24 hours later. Representative spectral cytometry plots at 24 hr showing *Il10*-eGFP against CD4 in *Nr4a3*-Timer^+^ **(L)**. Summary of *Il10*-eGFP^+^ frequency within the CD4^+^*Nr4a3*-Timer^+^ T-cells. **(M).** Analysis for additive effect of treatment by factoring αIFNγ and αIL-27 as two vairables **(N)**. **(C, D, G, I-K, M, N)** bars represent median with interquartile range. Statistical analysis by Mann-Whitney U Test (**C,D,I-K**), Kruskal-Wallis test with Dunn’s multiple comparisons (M). N = 4 **(A-E)**, N = 8 **(F-K)**, N = 15 (Combo), N = 16 (Isotype) N = 17 (αIFNγ) and N = 17 (αIL-27) **(L-N)** per treatment.

We were interested to understand relationships between NK cells and the development of Tr1 cells (identified as *Il10-*eGFP^+^ T-cells). CD4^+^ Timer^+^ *Il10*-eGFP^+^ T-cells directly correlate with the NK1.1^+^ TCR^−^ population under isotype treatment, but αIFNγ decouples this, removing the correlation between them (**Figure 4E**). Based on this, we hypothesised that NK1.1^+^ cells, as major *Ifng*-YFP expressors, may be the important physiological source of IFNγ that contributes to the regulation of Tr1 cell development. To determine the functional role of NK cells, we performed antibody depletion of NK1.1 cells before immunisation with [4Y]-MBP to induce Tr1 cells *in vivo*. Spectral cytometry confirmed staining for NK1.1^+^ (using a different antibody clone) that very few NK1.1^+^ cells remained following administration of the depleting antibody (**Figure 4F-G**). However, NK cell depletion had no effect on the frequency of *Il10-*eGFP^+^ T-cells (**Figure 4H-I**) nor on markers of strong TCR signalling ICOS (**Figure 4J**) or OX40 (**Figure 4K**). These data therefore suggest that NK cell IFNγ is redundant for Tr1 cell development in this model and indicating that T-cell derived IFNγ is the more important source.

### IFNγ and IL-27 additively regulate Tr1 cell development

Interleukin-27 (IL-27) is a potent, innate immune cell-derived inducer of IL-10 production ^11^. We were interested to compare the relative potency of IL-27 versus IFNγ for the induction of Tr1 cells in our accelerated adaptive tolerance model. We administered αIFNγ and/or αIL-27 along with immunisation of Tg4 *Nr4a3*-Tocky *Il10*-eGFP mice with 4 mg/kg [4Y]-MBP to examine the effect on Tr1 cell development (**Figure 4L-M**). In this setting, co-blockade of IL-27 and IFNγ resulted in a highly significant reduction in Tr1 cell frequency (**Figure 4M**), and analysis showed that this affect appeared to be additive when visualising αIL-27 and αIFNγ treatment as factored variables (**Figure 4N**).

## Discussion

Negative feedback control of T-cell responses by IL-10 is important in regulating Th1 cells that have undergone repeated stimulation ^8^. In our study we have demonstrated that the Th1-associated cytokine IFNγ indirectly promotes *Il10* expression in Tr1 cells through enhancing TCR signalling, highlighting a feedback loop that may act to control immunopathology. IL-10 increased IFNγ and FasL in TCR-stimulated CD8^+^ T-cells *in vitro* ^22^. However, it also activates Natural Killer cells with increased cytotoxic function ^23,24^. Despite this, IL-10 has been known to inhibit IFNγ production by NK cells in the presence of APCs, partially due to a decrease in IFNγ-inducing cytokines ^3,25^. This observation is consistent with our finding that under conditions of αIFNγ treatment, which lowers Tr1 cell frequency, we saw a concomitant increase in NK cell IFNγ.

Thymic FoxP3^+^ Treg comprise a self-reactive polyclonal T-cell population that undergo thymic selection in response to strong and persisting TCR signalling ^26,27^. Likewise, Tr1 cells have been previously reported to progressively develop by strong ^19^ and/or chronic ^16,17^ antigenic stimulation in the periphery ^9^. *Il10*^+^ T-cells induced by strong TCR signalling in our accelerated adaptive tolerance model are highly refractory to re-stimulation ^19^, suggesting IL-10 marks T-cells in an acute state of negative feedback. Mechanistically this has been linked to chromatin remodelling of regulatory gene regions in tolerogenic T-cells, whereby anti-inflammatory regions remain accessible to sub-threshold TCR signals whilst inflammatory gene loci are silenced ^28^. Induced Tr1 cells and peripheral FoxP3^+^ Treg may share some evolutionary redundant functions, since genetic deletion of the conserved non-coding DNA sequence one (CNS1) of the *Foxp3* gene leads to loss of FoxP3^+^ T-cells within the mesenteric lymph node but is compensated for by an expansion of Tr1-like IL-10^+^ T-cells ^29^. This suggests that IL-10^+^ T-cells subsets may be able to compensate for each other in some circumstances, e.g., at environmental interfaces where IL-10 is critical ^30^.

IL-10 is elicited preferentially in Tr1 cells when they are strongly stimulated through their TCR under tolerogenic conditions ^28^. Clearly, however, TCR signalling alone is unlikely to be the sole promoting factor for IL-10 expression and other signals are likely to contribute. Tolerogenic stimulation, however, is likely to provide a different activation pattern of nuclear transcription factors than T-cells activated in the context of enhanced co-stimulatory molecule expression. Many features of so called “exhausted” T-cells or Tr1 cells have been linked to chromatin remodelling ^28,31^. The relative activity of nuclear factor of activated T-cells (NFAT) alone versus NFAT:AP-1 complexes is thought to be important in co-ordinating the dysfunctional T-cell programme ^28,31^ and may it will be interesting to explore whether similar mechanisms are involved in the differentiation of Tr1-like cells under tolerising conditions.

It is intriguing that *Il10* can be rapidly (within four to twelve hours) transcribed in response to a single dose of soluble peptide and suggests that the *Il10* locus can be rapidly remodelled within the CD4^+^ T-cell population. As to whether this is unique to the subset transcribing *Il10* or is a general feature of a population of T-cells that have been strongly stimulated through their TCR remains to be determined. It also raises the question as to whether further heterogeneity exists within the CD4^+^ T-cell population that may pre-dispose some populations to *Il10* transcription over others (e.g., potential memory-like subsets or T-cells experiencing different grades of tonic signal in different environments ^32^). It is also clear from our data that aspects of the Tr1 transcriptional signature generated through our model are dynamic with some signature genes transcriptionally dampened 24 hr after the primary immunisation ^19^. Indeed, this may be because Tr1 phenotypes reflect CD4^+^ T-cells that have undergone recent and relatively strong TCR signalling in the absence of major co-stimulation and are in an acute state of negative feedback.

Our findings that macrophage activation status, directly correlated with the emergence of IL10^+^ T-cell expression suggests that, aside from their increased antigen-presenting capacity due to IFNγ priming, they may also provide additional signals. For this reason, we explored the potential role of IL-27 in our model. IFNγ and MYD88 are known to upregulate IL-27 within macrophages ^33^, so it is possible that IFNγ may work through both augmentation of TCR signal strength and the induction of the Tr1-promoting cytokine IL-27. It remains to be determined how the inclusion of innate pathogen recognition receptor signals would alter the differentiation of T-cells experiencing strong TCR signalling within the immune synapse. In addition, activation of APCs would lead to production of T helper cell polarising cytokines that may influence the activation of transcription factors to promote alternative T helper cell differentiation pathways, such as the Th1 pathway. We observed a small additive effect when we neutralised both IL-27 and IFNγ (almost halving the development of Tr1 cells *in vivo*), which suggests their mechanisms of action may not be entirely overlapping.

It is well established that IL-10 expression is a signature cytokine of the effector Treg programme ^34^, which is initiated by TCR signalling in Foxp3^+^ Treg cells ^35^. This strongly suggests that the *Il10* region is rapidly responsive to TCR stimulation in differentiated Treg, and *in vivo* αCD3 stimulation can enhance IL-10 expression *in vivo* during steady state ^36^. In a *T. gondii* infection model IFNγ and IL-27 have been shown to promote CXCR3 and T-bet in Tregs, but by separate mechanisms at separate sites – the periphery and musca respectively ^37^. They also found that Tregs exposed to either IFNγ or IL-27 have distinct transcriptional profiles. Whether the local concentrations of these cytokines may also skew the phenotype of Tr1 cells to promote their recruitment to different anatomical sites remains to be explored.

In summary, our study has established that Tr1 cells can be rapidly generated *in vivo* in response to strong tolerogenic signalling. Our data support that local IFNγ and IL-27 combine to additively regulate their generation in vivo, highlighting a new ‘regulatory’ role for IFNγ in the induction of tolerogenic Tr1 cells.

## Methods

### Mice

Tg4 *Nr4a3*-Tocky *Il10*-eGFP mice were used as previously described ^19^. *Nr4a3*-Tocky Tg4 Tiger (*Il10*-eGFP) were mated to Great (*Ifng*-YFP) Smart-17A^38^ mice to generate *Nr4a3*-Tocky Tg4 Tiger (*Il10*-eGFP) Great (*Ifng*-YFP) Smart-17A mice. *Nr4a3*-Tocky mice ^26^ were originally obtained under MTA from Dr Masahiro Ono, Imperial College London, UK. This manuscript does not report any findings arising from the use of unmodified *Nr4a3*-Tocky lines. All animal experiments were approved by the local animal welfare and ethical review body and authorised under the authority of Home Office licenses P18A892E0A and PP3965017 (held by D.B.). Animals were housed in specific pathogen-free conditions. Both male and female mice were used, and littermates of the same sex were randomly assigned to experimental groups.

### Immunisations

Tg4 *Nr4a3*-Tocky Tiger (*Il10*-eGFP) mice were immunized through subcutaneous injection of [4Y]-MBP peptide (MBP Ac1-9[4Y] peptide AcASQYRPSQR, GL Biochem, Shanghai) with doses stated in figure legends in a total volume of 200 μL PBS. Mice were then euthanised at the indicated time points, and spleens removed to analyse systemic T-cell responses.

### Antibody treatments

For *in vivo* blockade experiments, *in vivo* grade anti-IL-27p28 (clone MM27.7B1, mouse IgG2a, BioXcell), or anti-IFNγ (clone XMG1.2, kind gift from Prof Anne Cooke, University of Cambridge, rat IgG1) were administered through intraperitoneal injection as indicated in figure legends. For anti-IFNγ experiments, an isotype control group was used consisting of rat IgG1 (clone MAC221, kind gift from Prof Anne Cooke, University of Cambridge) and for anti-IL-27 experiments, an isotype control group was used consisting of mouse IgG2a (clone C1.18.4, BioXCell). For NK cell depletion, 200 µg of depleting NK1.1 (clone PK136, BioXCell) or mouse IgG2a (clone C1.18.4, BioXCell) was administered 48 hours before peptide immunisation.

### Flow cytometry and cell sorting

For analysis of splenic lymphocytes single cell suspensions were prepared as previously described ^39^. For analysis of splenic myeloid populations spleens were dissociated using scissors in 1.2 mL of digestion media containing 1 mg/mL collagenase D (Merck Life Sciences) and 0.1 mg/mL DNase I (Merck Life Sciences) in 1 % FBS (v/v) RPMI. Samples were then incubated for 20-25 min at 37 °C in a thermo-shaker. Digestion mixture was then passed through a 70 µm filter (BD Biosciences) and washed with 30 mL ice cold media (10 % FBS RPMI). Digested cells were washed once and stained in 96-well U-bottom plates (Corning). Analysis was performed on a BD LSR Fortessa X-20 instrument. The blue form of the Timer protein was detected in the blue (450/40 nm) channel excited off the 405 nm laser. The red form of the Timer protein was detected in the mCherry (610/20 nm) channel excited off the 561 nm laser. A fixable eFluor780-flurescent viability dye (eBioscience) was used for all experiments. Directly conjugated antibodies used in these experiments were: From BioLegend: Rat Anti-Mouse CD19 PerCP-Cy5.5 (clone 6D5), Rat Anti-Mouse CD25 PerCP-Cy5.5 (clone PC61), Rat Anti-Mouse CD40 PerCP-Cy5.5 (clone 3/23), Rat Anti-Mouse OX40 PerCP-Cy5.5 (clone OX-86), Rat Anti-Mouse PD-1 PerCP-Cy5.5 (clone 29F.1A12), Rat Anti-Mouse CD11b APC (clone M1/70), Armenian Hamster Anti-Mouse CD11c APC (clone N418), Rat Anti-Mouse CD25 APC (clone PC61), Armenian Hamster Anti-Mouse CD69 APC (clone H1.2F3), Rat Anti-Mouse LAG3 APC (clone C9B7W), Rat Anti-Mouse OX40 APC (clone OX-86), Rat Anti-Mouse PD-1 APC (clone 29F.1A12), Rat Anti-Mouse PDL1 APC (clone 10F.9G2), Rat Anti-Mouse/Human CD11b PE-Cy7 (Clone M1/70), Armenian Hamster Anti-Mouse CD11c PE-Cy7 (Clone N418), Rat Anti-Mouse CD19 PE-Cy7 (clone 6D5), Rat Anti-Mouse CD25 PE-Cy7 (clone PC61), Rat Anti-Mouse CD86 PE-Cy7 (clone GL-1), Rat Anti-Mouse ICOS PE-Cy7 (clone 7E.17G9), Rat Anti-Mouse LAG3 PE-Cy7 (clone C9B7W), Rat Anti-Mouse MHCII I-A/I-E PE-Cy7 (clone M5/114.15.2), Mouse Anti-Mouse NK1.1 PE-Cy7 (clone S17016D), Mouse Anti-Mouse TIGIT PE-Cy7 (clone 1G9), Rat Anti-Mouse CD4 AF700 (clone RM4-4), Armenian Hamster Anti-Mouse CD69 AF700 (clone H1.2F3), Armenian Hamster Anti-Mouse ICOS AF700 (clone C398.4A), Rat Anti-Mouse MHCII I-A/I-E AF700 (clone M5/114.15.2), Armenian Hamster Anti-Mouse TCRbeta AF700 (clone H57-597); From BD Biosciences: Armenian Hamster Anti-Mouse CD11c BUV395 (clone HL3), Rat Anti-Mouse CD4 BUV395 (clone GK1.5), Rat Anti-Mouse F4/80 BUV395 (clone T45-2342), Mouse Anti-Mouse TCRVbeta8.1/8.2 BUV395 (clone MR5-2), Rat Anti-Mouse CD11b BUV737 (clone M1/70), Rat Anti-Mouse CD19 BUV737 (clone 1D3), Rat Anti-Mouse CD4 BUV737 (clone GK1.5). For intracellular staining of Foxp3, the Foxp3 transcription factor staining buffer kit was used (eBioscience), using Rat Anti-Mouse FoxP3 APC (clone FJK-16s). For cell sorting, single cell suspensions from biological replicate mice were generated. Cells were sorted on a FACS ARIA FUSION cell sorter.

### Spectral cytometry

Cells were prepared as above in Flow cytometry and cell sorting, except the cells were stained in Brilliant Stain Buffer (BD Biosciences) and analysis was performed on a Sony ID7000 spectral cytometer.

### RNA-seq analysis

Data from GEO: GSE165817 (Elliot et al. ^19^) were re-analysed using DESeq2 ^40^ in R version 4.0. Normalized read counts were transformed using the regularised log (rlog) transformation. For transcriptional profiling of *Il10*-eGFP cells, RNA was extracted from lysates using the Arcturus Picopure RNA kit (ThermoFisher) according to the manufacturer’s instructions. 5 ng of RNA was used for generation of sequencing libraries using the Quantseq 3’ mRNA-seq Library Preparation kit FWD (Lexogen). Unique Molecular Identifiers (UMIs) were used for the evaluation of input and PCR duplicates and to eliminate amplification bias. Libraries were normalised and pooled at a concentration of 4 nM for sequencing. Libraries were sequenced using the NextSeq 500 using a Mid 150v2.5 flow cell. Cluster generation and sequencing was then performed and FASTQ files generated. FASTQ files were analysed using the BlueBee QuantSeq FWD pipeline and aligned to the GRCm38 (mm10) genome. HTSeq-count v0.6.0 was used to generate read counts for mRNA species and mapping statistics. Raw read counts in the .txt format were used for further analysis using DESeq2in R version 4.0 ^40^. A DESeq dataset was created from a matrix of raw read count data. Data were filtered to remove genes with fewer than 10 reads across all samples and one biological replicate pair was removed from analysis due to being identified as an outlier using principal component analysis. Heatmap analysis was performed on the rlog transformed data using the R package gplots.

### Statistical analysis

Statistical analysis was performed on Prism 10 (GraphPad) software. For comparison of non-parametric data, a Mann Whitney U test was performed or a Kruskal Wallis test with Dunn’s multiple comparisons test. For parametric data a student t test was used. For experiments with two factors, a two way ANOVA was performed with Sidak’s multiple comparisons testing. Variance is reported as median ± interquartile range or bars reflect the median for non-parametric data (unless otherwise stated); data points typically represent individual mice. *p = < 0.05, **p = < 0.01, ***p = < 0.001, ****p = < 0.0001.

## Acknowledgements

Work funded by the Wellcome Trust MIDAS 4-year PhD programme (D.A.J.L), Wellcome Trust Seed Award in Science (214018/Z/18/Z, D.B.), the Lister Institute of Preventative medicine (D.B., S.R.) and MRC Career development awards (MR/V009052/1, D.B., L.S.; MR/S024611, R.A.D.). L.S.G and D.C.W are funded by the University of Birmingham. We thank the Tech Hub flow cytometry team led by Dr Guillaume Desanti at University of Birmingham.

## Supplementary Figures

**Supplementary Figure 1: Tr1-like markers arise after early Th1 hallmarks**

Re-analysis of GEO: GSE165817 (Elliot et al. ^19^) where Tg4 *Nr4a3*-Tocky *Il10*-eGFP mice were administered with 0 (control), 8, or 80 μg [4Y]-MBP in 200 μL PBS s.c. and spleens harvested at 4, 12 and 24 hr for bulk RNA isolation and transcriptome analysis. Heatmap of markers associated with early T-cell activation, Th1 phenotype or Tr1-like cells arising after expression of Th1 cell hallmarks.

**Supplementary Figure 2: Effect of αIFNγ on B cell and Dendritic cell responses**

Tg4 *Nr4a3*-Tocky *Il10*-eGFP *Ifng*-YFP mice were administered 4 mg/kg [4Y]-MBP in PBS s.c. and 1 mg αIFNγ or αIgG1 isotype in 200 μL PBS i.p. and spleens were harvested. Representative flow plots at 24 hr showing MHCII against CD19 and MHCII^+^ CD19^−^ derived CD11c^±^ under αIFNγ or αIgG1 treatment **(A)**. Summary of changes in MHCII^+^ CD19^−^ **(B)**, MHCII^+^ CD19^+^ **(C)** from **(A)**, and CD11C^+^ **(D)** frequency from **(B)**. Summary of changes in PD-L1 MFI **(E)** from **(D). (B-E)** bars represent median with interquartile range. Statistical analysis by Mann-Whitney test. N = 8 **(B-E)** per treatment.

