## Supplementary figures and images for "Interferon-γ and IL-27 positively regulate type 1 regulatory T-cell development during adaptive tolerance"

### Supplementary Figures 1 and 2

#### Supplementary Figure 1

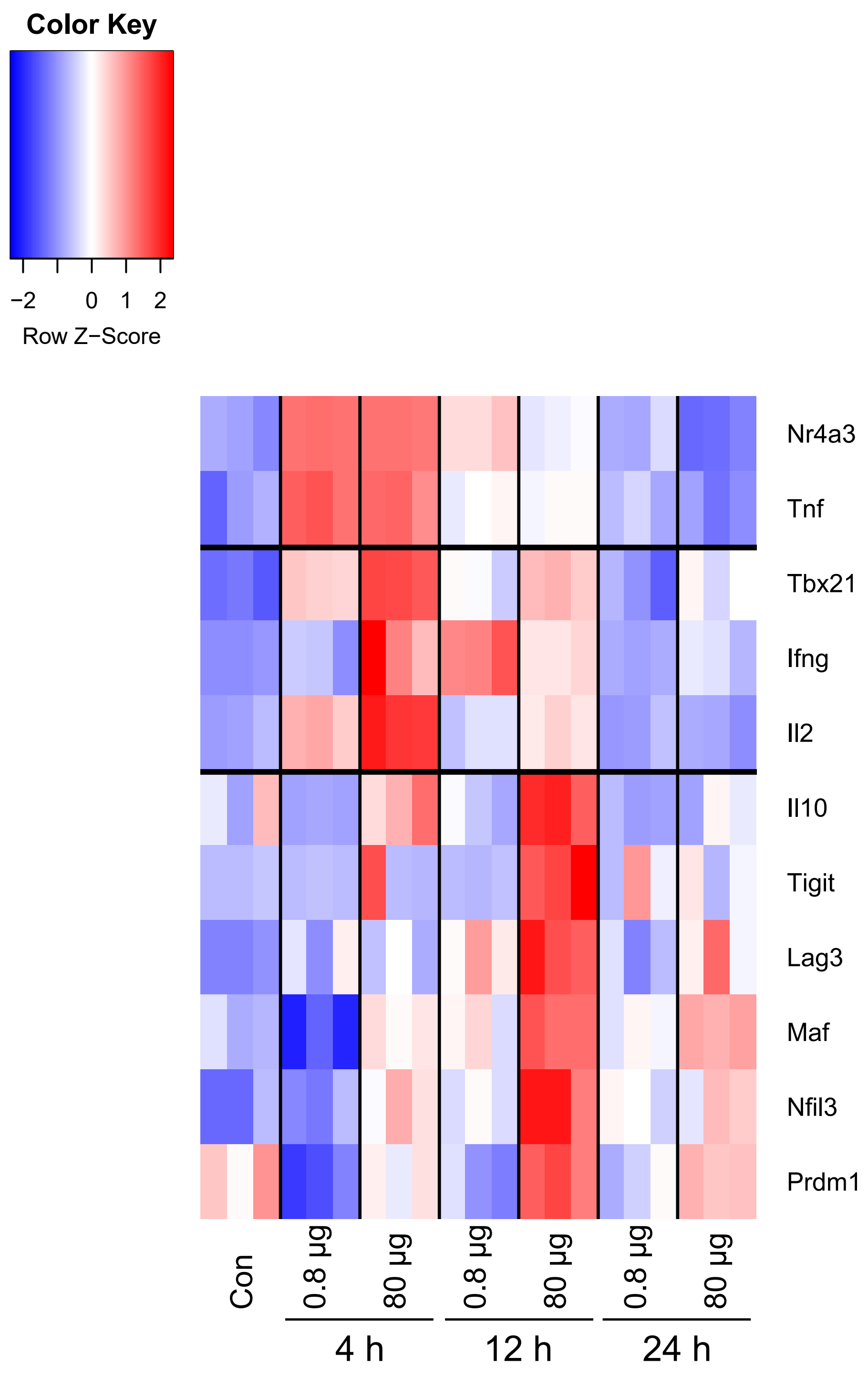

Supplementary Figure 2

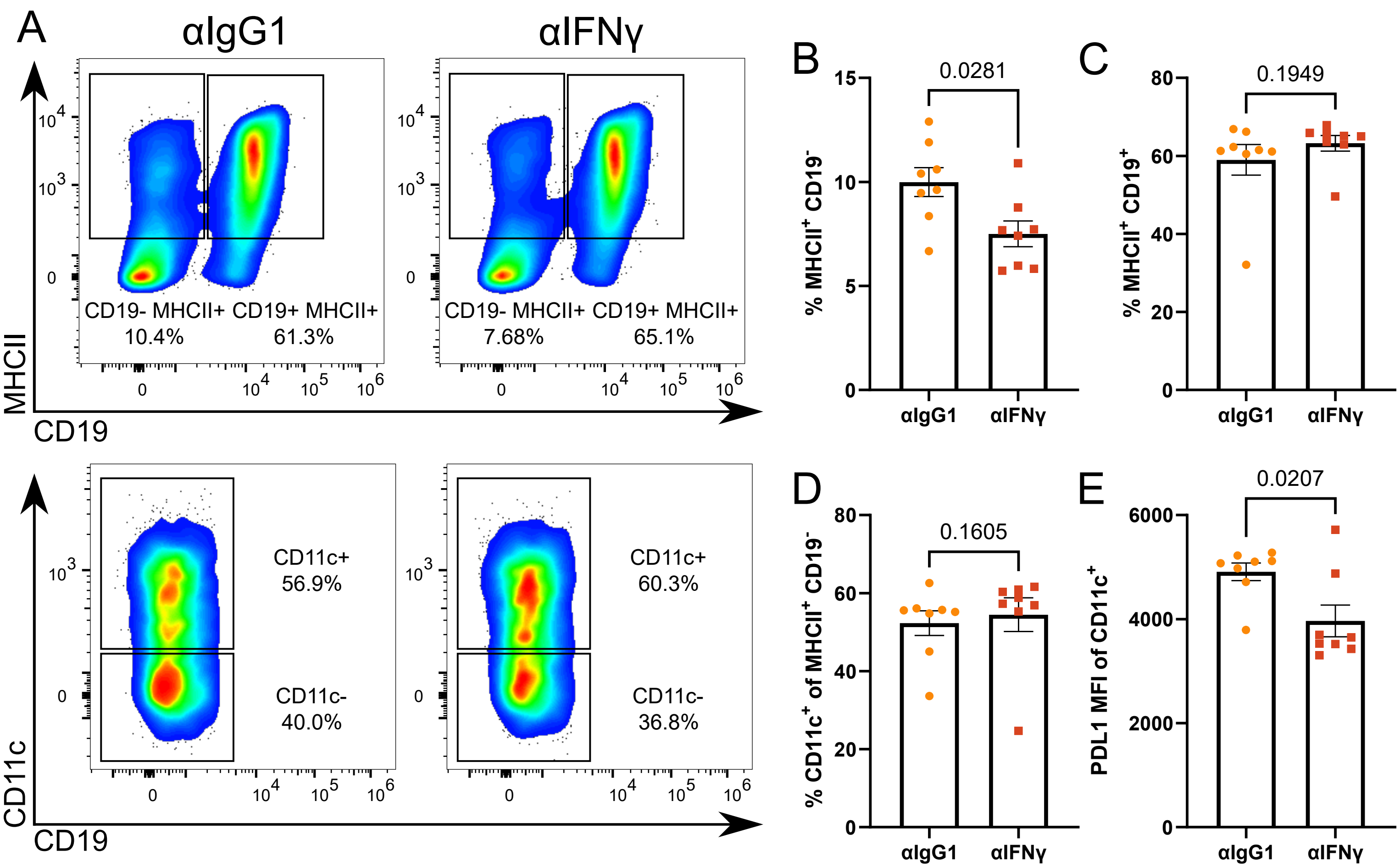
